# Asymptotically optimal minimizers schemes

**DOI:** 10.1101/256156

**Authors:** Guillaume Marçais, Dan DeBlasio, Carl Kingsford

## Abstract

**Motivation:** The minimizers technique is a method to sample *k*-mers that is used in many bioinformatics software to reduce computation, memory usage and run time. The number of applications using minimizers keeps on growing steadily. Despite its many uses, the theoretical understanding of minimizers is still very limited. In many applications, selecting as few *k*-mers as possible (i.e. having a low *density*) is beneficial. The density is highly dependent on the choice of the order on the *k*-mers. Different applications use different orders, but none of these orders are optimal. A better understanding of minimizers schemes, and the related local and forward schemes, will allow designing schemes with lower density, and thereby making existing and future bioinformatics tools even more efficient.

**Results:** From the analysis of the asymptotic behavior of minimizers, forward and local schemes, we show that the previously believed lower bound on minimizers schemes does not hold, and that schemes with density lower than thought possible actually exist. The proof is constructive and leads to an efficient algorithm to compare *k*-mers. These orders are the first known orders that are asymptotically optimal. Additionally, we give improved bounds on the density achievable by the 3 type of schemes.

## 1 Introduction

The *minimizers* technique is a method (Roberts *et al.*, 2004b,a; Schleimer *et al.*, 2003) to sample *k*-mers from a sequence. It has two important properties: (i) there is no large gap in the sampling and (ii) from similar sequences similar *k*-mers are sampled. Minimizers help design algorithms that are more efficient both in memory usage and run time by reducing the amount of information to process, while not losing information.

The minimizers method is very flexible and has been used in a surprising large number of settings, from the original computation of read overlaps (Roberts *et al.*, 2004b,a), to counting *k*-mers (Li and XifengYan, 2015; Deorowicz *et al.*, 2015), reducing the genome assembly de Bruijn graph (Ye *et al.*, 2012; Li *et al.*, 2013), making sequence alignment faster (Li, 2016; Ondov *et al.*, 2016), to metagenomics (Wood and Salzberg, 2014; Kawulok and Deorowicz, 2015) and sparse data structures (Grabowski and Raniszewski, 2015).

The minimizers method has two parameters, *k* the length of the *k*-mers and *w* the maximum distance between two sampled *k*-mers in the input sequence (called the window size). Additionally, the minimizers scheme is parameterized by the choice of a complete order on the *k*-mers, for example the lexicographic order. The minimizers scheme then selects in the input sequence the smallest *k*-mer, according to the predefined order, in each window of *w* consecutive *k*-mers.

Any choice of an order on the *k*-mers is valid in the sense that properties (i) and (ii) above are satisfied. The *density*, i.e. the expected number of selected *k*-mers over the length of the input sequence, is affected by the choice of the order on the *k*-mers. For many applications, a lower density is preferable as it reduces the amount of data to process. The density of any minimizers scheme is at least 1*/w*, as at least 1 *k*-mer is selected in each window, and at most 1, when every *k*-mer in the sequence is selected. These are the trivial bounds.

Although bioinformatics tool developers chose many different orders for their applications, in most cases, these orders are not an integral part of their algorithm. That is, if one were to change the order in a given application, the results returned by the application would be unchanged or equivalent, although the run time or memory usage may be different. Therefore, the development of *k*-mer orders giving lower density would benefit new and existing applications.

In addition to minimizers schemes, we consider two generalizations: local schemes and forward schemes. Local schemes are the most general schemes as they use any function defined on a set of *w k*-mers. The forward schemes are local schemes with the additional requirement that the *k*-mers are selected in an increasing manner from the input sequence (minimizers schemes are an example of forward schemes). From the more specific to the more general schemes, we have

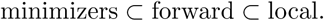

Similarly, from the point of view of a bioinformatics application, these other two types of schemes can be used as a drop-in replacement of a minimizers scheme: they have the same parameters *k* and *w*, the same input and output, and they satisfy properties (i) and (ii).

Schleimer *et al.* (2003) showed an expected density of 2*/*(*w* + 1) for some class of minimizers schemes and proposed a lower-bound for local schemes of (1.5 + 1/2*w*)/(*w* + 1). Marçais *et al.* (2017) used a heuristic method to create minimizers schemes with density below 2/(*w* + 1), but still above that lower bound for local schemes.

We study the asymptotic behavior in *k* and *w* of local, forward and minimizers schemes. This study leads to the realization that the previously believed lowerbound on the achievable density of local schemes does not apply. We give concrete examples of minimizers schemes with density below that former lower-bound. This opens up the field for much greater improvements in density from minimizers, forward and local schemes. Also, the results of this study gives directions for further potential improvements.

We present three main results in this paper.

I. For fixed *w* and asymptotically in *k*, we show that there exist minimizers schemes that achieve optimal density. This is the first example of schemes that achieve optimal density. Because minimizers schemes are the most specific schemes, optimality is also achieved by local and forward schemes, and this completely characterizes the behavior for fixed *w* and asymptotic in *k*.
II. For fixed *k* and asymptotically in *w*, we derive new lower-bounds for minimizers and forward schemes, which show that neither of these schemes can be optimal when *w* is much larger than *k*. Furthermore, we show that the local schemes are strictly more powerful than forward schemes by finding optimal local schemes for some parameters *k* and *w* with density not achievable by forward schemes.
III. We prove a lower-bound on the density of forward schemes that is valid for all parameters *k* and *w*. This lower-bound is a refinement of the bound proposed by Schleimer *et al*. (2003).

Additionally, these results give two practical algorithms. The first one computes a set of *k*-mers that covers every path of length *w* in the de Bruijn graph (an extension of the set cover problem). This problem was studied in Orenstein *et al*. (2017) and Marçais *et al*. (2017), and this new algorithm gives an asymptotically optimal solution. The second algorithm gives the order between *k*-mers for the minimizers schemes in (I). This algorithm is efficient as it only takes *O*(*k*) time to compare two *k*-mers.

In the next section, we give precise definitions of the various concepts and a statement of the main theorems. Section 3 gives details proofs of the theorems, followed by a discussion of remaining open problems (section 4).

## 2 Approach

### 2.1 Definitions

#### Basic definitions

Let Σ be an alphabet of size *σ* = |Σ|. If *S* ∈ Σ^*^ is a string on alphabet Σ, *S*[*i*, *ℓ*] is the substring of *S* starting at position *i* and of length *£*. In many cases in the following the strings are circular: offsets in the string are understood modulo the length of the string and a substring extending beyond the end of the string wraps around to the beginning of the string. The substring *ω* = *S*[*i*, *w* + *k* − 1] represents the sequence of a window *ω* of *w* consecutive *k*-mers starting at offset *i* in *S*.

#### Schemes

A *local scheme* is a function that selects a *k*-mer in a window of *w* consecutive *k*-mers, i.e. *f*: Σ*^w+k−1^* → [0: *w* − 1].

A *forward scheme* is a particular local scheme where the sequence of the starting positions of the selected *k*-mers is an increasing sequence. Equivalently, a local scheme *f*: Σ*^w+k−1^ →* [0: *w* − 1] is a forward scheme if

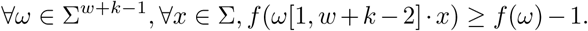

The dot is the concatenation operator and *ω*[1, *w* + *k* − 2] · *x*, ∀*x* ∈ Σ represents all the possible windows following *ω*.

A *minimizers scheme* is a particular local scheme where the function returns the left-most position of the smallest *k*-mer in the window. All minimizers schemes are forward schemes, and forward schemes are local schemes, but those sets are not equal to each other.

#### Density

The set of *selected indices* of a scheme *f* on string *S* is

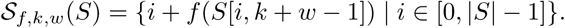

Because the scheme may select in adjacent windows the same position in *S*, 
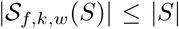
. The *particular density* of a scheme *f* on the circular string *S* is the proportion of *k*-mers selected: 
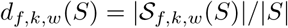
.

The *density* of a scheme *d_f,k,w_* is defined at the limit as the expected density on an infinitely long sequence with the characters selected IID (Independent Identical Distribution). The trivial bounds for the density are 1/*w* ≤ *d_f,k,w_* ≤ 1. Following Marçais *et al*. (2017), we define the *density factor* df = *d* · (*w* + 1), which represents the average number of *k*-mers selected in every window of length *w* + 1. The trivial bounds for the density factors are 1 + 1/*w* ≤ df*_f,k,w_* ≤ *w* + 1.

#### Computing density

A *de Bruijn sequence* of order *ℓ* is any circular sequence of length *σ*^*ℓ*^ such that every possible substring of length *ℓ* occurs once and only once in the string (de Bruijn, 1946). In the following, *D*_*ℓ*_ represents the de Bruijn graph of order *ℓ*. A de Bruijn sequence is obtained by reading the first base of every vertex traversed by a Hamiltonian tour of a de Bruijn graph *D*_*ℓ*_ of order *ℓ*.

Even though the density is defined as the limit of an expected value on an infinite string, it can be computed exactly (and not just estimated) as the particular density computed on a de Bruijn sequence of large enough order (Marçais *et al*., 2017). For any de Bruijn sequence *S*_*ℓ*_ of order *ℓ* ≥ 2*w* + *k* − 2, the particular density of *f* on *S*_*ℓ*_ is equal to the density of *f*: *d_f,k,w_* = *d_f,k,w_* (*S*_*ℓ*_). For a forward scheme, the minimum order of the de Bruijn sequence is only *w* + *k*.

#### Universal set

A *universal set* is an unavoidable set of *k*-mers: it is a set *U_k,w_* such that every path of *w* nodes in the de Bruijn graph of order *k* contains a *k*-mer from *U_k,w_*. In other words, a universal set is a set of nodes of *D_k_* that covers every path of *w* nodes. Equivalently, every string of length *k* + *w* − 1 must contain a substring of length *k* from *U_k,w_*.

There is a strong link between universal sets and minimizers schemes. A minimizers scheme is *compatible* with a universal set *U_k,w_* if every *k*-mer of *U_k,w_* compares less than any *k*-mer not in *U_k,w_*. There is more than one order which is compatible with a universal set, as the relative order of the *k*-mers within *U_k,w_* is not constrained by the definition. Although this relative order may change the density of the scheme, it is not relevant in our asymptotic analysis.

### 2.2 Main results

#### Behavior asymptotically in *k*

We show that for any fixed value of *w*, there exists a sequence of minimizers schemes that asymptotically achieve optimal density, i.e. one *k*-mer per window or 1/*w*. These are the first proposed orders that achieve close to optimal density. It is also the first orders to have density factors below 1.5 + (1/2*w*), which was formerly considered the lowest possible density factor. Surprisingly, this optimal density is attained with minimizers schemes, the weakest type of schemes.

The sequence of minimizers schemes is created from universal sets. The following Lemma shows an important link between universal sets and minimizers scheme: the size of a universal set upper-bounds the density of any compatible minimizers scheme.

##### Lemma 1.

Given a minimizers scheme *f_U_* compatible with a universal set *U*, the density satisfies

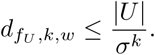

The strategy is then to construct a sequence of universal sets whose sizes get close to *σ^k^*/*w*. This gives us our first main result, that optimal minimizers schemes exist asymptotically in *k* for alphabet of even sizes.

##### Theorem 1.

On an even alphabet, for any fixed *w*, there exists a sequence of universal sets *U_k_* asymptotically of optimal size, and a sequence of minimizers schemes *f_k_* asymptotically of optimal density.

The proof of this theorem relies on a geometric argument. The de Bruijn graph is embedded in a *w*-dimensional space, a hypercube with *w* dimensions, such that the *k*-mers mapping into a particular volume of the cube is a universal set. Then, we show that asymptotically the number of *k*-mers mapping into that volume represents a proportion 1/*w* of the total number of *k*-mers. This line of proof is a generalization of the construction of asymptotically minimum vertex cover by Lichiardopol (2006).

#### Behavior asymptotically in *w*

The previously mentioned bounds on density of 2/(*w* + 1) might give the impression that, as *w* gets large, arbitrarily small density can be achieved. It is not the case for minimizers schemes which have a lower limit on the density greater than 0.

##### Theorem 2.

For any minimizers scheme *f*, the density *d_f,k,w_* converges asymptotically in *w* to *σ^−k^*:

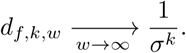

Forward schemes and local schemes do not have such a lower-bound on their density, and we will construct a forward scheme with a density going to 0 asymptotically in *w*. In such cases, it is more meaningful to speak of the density factor. The lowest density factor for minimizers is Ω(*w*), while the lowest density factor for forward schemes is 
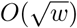
.

#### Forward schemes bound

Finally, we prove a lower-bound on the density for forward schemes, which holds for any parameters *k* and *w*.

##### Theorem 3.

The density of any forward scheme satisfies

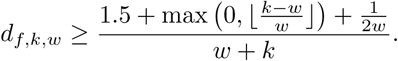

Asymptotically in *w*, this bound implies that the best density factor for forward schemes is ≥ 1.5. This is much lower than the lower-bound for minimizers schemes, although it is not known yet how to construct a forward scheme approaching that lower-bound for large values of *w*.

## 3 Proofs of main theorems

### 3.1 Minimizers asymptotic behavior in *k*

We consider in this section the behavior of minimizers schemes when the length of the window *w* is fixed while the length of the *k*-mer goes to infinity. In particular, we will construct asymptotically optimal minimizers schemes. Given that a local scheme must select at least one *k*-mer in each window, the minimum density is ≥ 1/*w*. So, more precisely, we construct a minimizers scheme *f* such that 
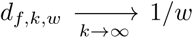
. To do so, we use the link between universal sets and orderings.

Recall that given a universal set *U*, a set that intersects every *w*-long path in the de Bruijn graph *D_k,σ_*, the minimizers scheme *f* is compatible with *U* if every *k*-mer of *U* compares less than any *k*-mer not in *U*. The following Lemma explains the fundamental link between universal sets and orderings.

#### Lemma 1.

Given a minimizers scheme *f_U_* compatible with a universal set *U*, the density satisfies

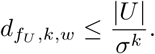

##### Proof.

Let *S_l_* be a de Bruijn sequence of order *ℓ* ≥ *w* + *k*. Because *U* is a universal set, every window of *w* consecutive *k*-mers contains at least one element of *U*, hence all the selected *k*-mers are from the set *U*. Therefore, 
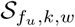
 contains, at most, all the positions in *S* of the *k*-mers of *U*. Moreover, every *k*-mer occurs exactly *σ^l−k^* times in *S_l_*. Hence

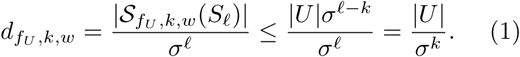

This Lemma gives a simple lower-bound on the size of a universal set for a de Bruijn graph. Define *β_w_* (*G*) to be the minimum size of a universal set for a graph *G* that hits every path of *w* vertices. This notation is an extension of the definition of the size of the minimum vertex cover: *β_2_*(*G*) = *β*(*G*).

#### Proposition 1.

The minimum size of a universal set satisfies:

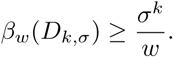

##### Proof.

Let *U* be a universal set of minimum size, |*U*| = *β_w_* (*D_k,σ_*), and *f_U_* a minimizers scheme compatible with *U*. By Lemma 1

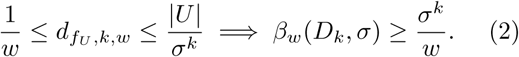

A second consequence of Lemma 1 is that a sequence of universal set *U_k_* that is asymptotically optimal, i.e. such that 
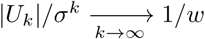
, gives a sequence of minimizers schemes *f_U_k__* which is also asymptotically optimal, i.e. 
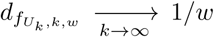
. The remainder of this section describes how to construct such a sequence of universal sets. This construction is similar to the construction of an optimal vertex cover (Lichiardopol, 2006) and of an optimal decycling set (Mykkeltveit, 1972).

#### 3.1.1 Naive extension

Given a universal set *U* that hits every *w*-long path in *D_k,σ_*, we can easily construct a universal set that hits every *w*-long path in *D_k+1,σ_*. The set *U* · Σ obtained by concatenating every letter of the alphabet Σ to every element of *U* is called the *naive extension*. The naive extension of a universal set *U* in *D_k,σ_* is a universal set in *D_k+1,σ_* for *w*-long paths, but the naive extension may not be of optimal size, even if *U* is of optimal size.

The naive extension shows that the minimum size of a universal set, in proportion, is a non-increasing function as the length of the *k*-mers increases.

##### Proposition 2.

*The function k → β_w_ (D_k,σ_)/σ^k^ is non-increasing*.

###### Proof.

Let *U_k,w_* be a universal set of minimum size, i.e. |*U_k,w_*| = *β_w_* (*D_k,σ_*). Consider now the naive extension *U_k,w_* · Σ. The size of the naive extension is |*U_k,w_* · Σ| = *σ*|*U_k,w_*|. Then,

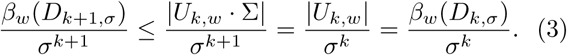

In the following construction, we only consider a subsequence of universal sets, where *k* is a multiple of *w*. Nevertheless, using the naive extension, the subsequence can be extended to a non-increasing complete sequence of universal sets that is asymptotically optimal.

#### 3.1.2 Embedding of the de Bruijn graph

The universal set is created by embedding the de Bruijn graph in a *w*-dimensional space and picking a region of the space that intersects every path of length *w*. Using a mapping (called *ψ_w_*), we embed the de Bruijn graph *D_k_* in a *w*-dimensional hypercube 
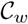
, where edges in *D_k_* correspond approximately to a rotation around a diagonal of the hypercube (a rotation plus a small translation). The hypercube is then split into *w* volumes, or wedges called *W_i_*, *i* ∈ [0, *w* − 1], of equal size, and one of them is selected, say *W_0_*. Because of the edges do not correspond perfectly to a rotation, the *k*-mers mapping to the wedge *W_0_* do not constitute a universal set. Therefore, we enlarge the wedge *W_0_* to get a universal set: *W̅_0_* is the wedge *W_0_* augmented with thin “slabs” at the frontier between *W_0_* and the other wedges. The set *W̅_0_* is such that any path of *w* vertices in *D_k_* contains at least one vertex that maps to *W̅_0_*. Finally, the universal set is any *k*-mer that maps into the wedge *W̅_0_*, that is the set 
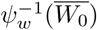
.

By construction, the same number of *k*-mers maps to any of the *w* wedges. The technical part of the proof below is to show that the number of *k*-mers mapping into the extra slabs is negligible as *k* → ∞.

In the following, we assume that *w* divides *k* and set *n* = *k*/*w*. Also, the alphabet is mapped to the integers {0,…, *σ* − 1}. For a *k*-mer *m*, define the functions

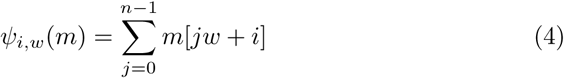

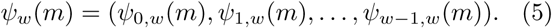

*ψ_i,w_* (*m*) sums every *w*th bases of *m*, starting at base *i*. *ψ_i,w_* (*m*) is an integer in the range [0: *n*(*σ* − 1)]. Then *ψ_w_* (*m*) is a *w*-dimensional vector that maps a *k*-mer to a point with integer coordinates inside a *w*-dimensional hypercube of side length *n*(*σ* − 1) + 1 (see Figure 1).

**Figure 1:**
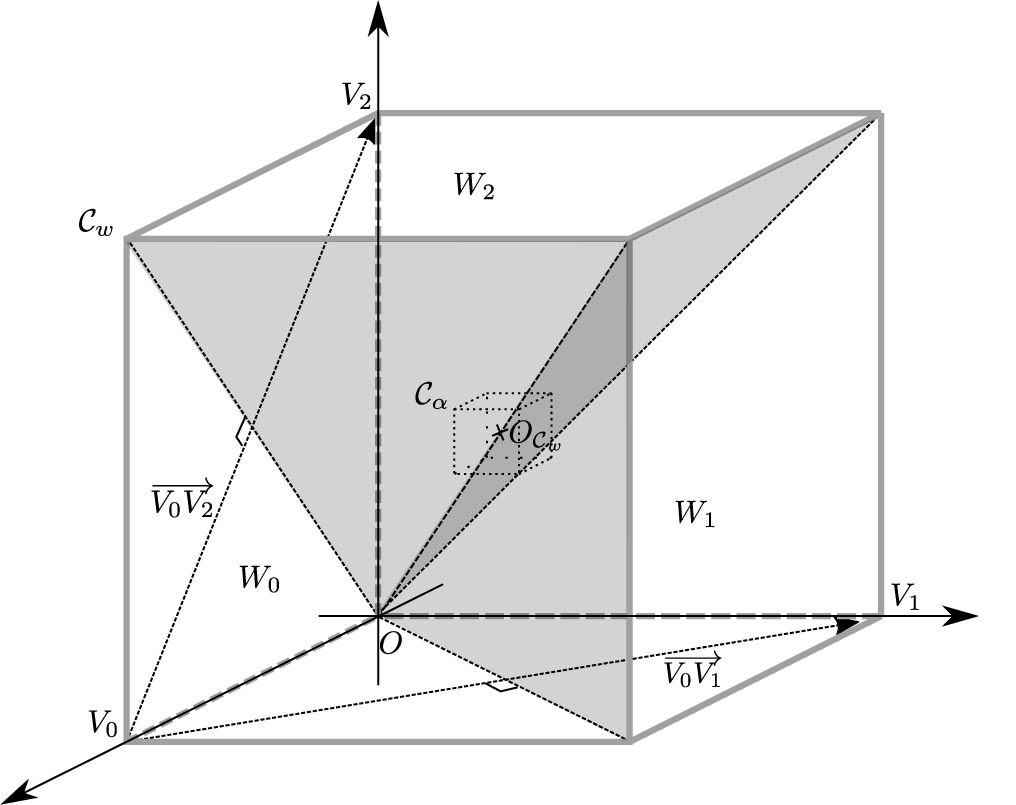
The cube 
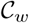
 for *w* = 3. The vertices *V_i_* have their *i*th coordinate equal to *n*(*σ* − 1), where *n* = *k*/*w*, and all other coordinates equal to 0. The wedge *W_i_* is the part of the cube of the points whose *i*th coordinate is greater than the others, which includes vertex *V_i_*. The hyperplanes separating separating wedge *W_i_* and *W_j_* is orthogonal to the vector *V*_*i*_*V*_*j*_. An edge in the de Bruijn graph corresponds to a rotation of 2*π*/*w* around the line 
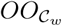
, followed by a translation along the (*w* − 1)th coordinate of at most *σ*.

The mapping *ψ_w_* has an important property: an edge in the de Bruijn graph corresponds to a rotation in the space, plus a translation along the last coordinate of length at most *σ* − 1. More precisely, given an edge *m* → *m_I_* in *D_k_*, the suffix of *m* is equal to the prefix of *m′*: *m′*[*i*] = *m*[*i* + 1], *i* ∈ [0, *k* − 2]. Hence, there is a shift in most of the coordinates in the mapping: *ψ_i,w_*(*m′*) = *ψ_i+1,w_* (*m*), *i* ∈ [0, *w* − 2]. Only the last coordinate of *ψ_w_* (*m′*) is not directly equal to a coordinate of *ψ_w_* (*m*), but rather *ψ_w−1,w_*(*m′*) = *ψ_0,w_* (*m*) − *m*[0] + *m′*[*k* − 1]. In other words, *ψ_w_* (*m′*) is obtained from *ψ_w_* (*m*) by rotating all the coordinates and adding a vector *δ* which has only one non-zero coordinate *δ*[*w* − 1] ∈ [−(*σ* − 1): *σ* − 1]:

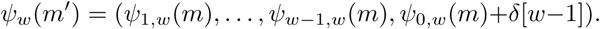

We define the wedge *W_i_* as the points in the hypercube whose *i*th coordinate is greater than all the other coordinates. Let *V_i_* be a vertex of the hypercube whose only non-zero coordinate is the *i*th coordinate. If *x* ∈ *W_0_*, then its 0th coordinate is greater than its *i*th coordinate: 
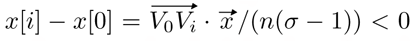
. The wedge *W_0_* is then equivalently defined as the volume bounded by the *w* − 1 hyperplanes orthogonal to the vectors 
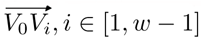
:

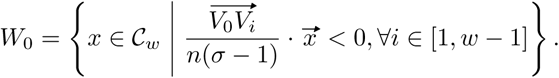

Notice that the hyperplanes separating the wedges are not contained in any of the wedges, and therefore the wedges are disjoint and have the same size.

The rotation of the coordinates correspond geometrically to a rotation around the vector (1,…, 1), and this vector is contained in the intersection of all the hyperplanes.

Ignoring at first the points on the hyperplanes, take a point *x* of 
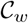
: it is in one of the wedges, say *W_i_*. The rotation of the coordinates of *x* gives a point in *W_i−1_*(or *W_w−1_* if *i* = 0). Hence, either *x* is in *W*_0_ or one of its *w* − 1 consecutive rotations is in *W_0_*. If it were not for the additional translation by *δ*, the *k*-mers mapping to *W_0_* would constitute a universal set.

Starting from a *k*-mer *m* that maps to point *x*, any *k*-mer reachable from *m* by a path of at most *w* − 1 edges maps to a point where each coordinate differs from a rotation of *x* by at most *σ*. To compensate for *δ* and the points on the hyperplanes, we extend the wedge *W_0_* by pushing the hyperplanes back by *σ*(i.e., we add some slabs parallel to the hyperplanes of thickness *σ*):

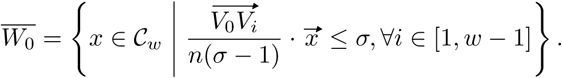

By construction, the set 
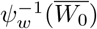
 is a universal set.

The slabs have a thickness of *σ*, which is a constant independent of *k*, hence only *w* − 1 dimensions of the slabs grow with *k* and the volume of the slabs is in proportion a negligible volume of the hypercube as *k* → ∞. The slabs are represented by the set *s* = *W̅*_0_\*W̅*_0_. It remains to show that the number of *k*-mers that map into the slabs, i.e. 
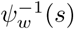
, is also asymptotically negligible compared to the total number of *k*-mers *σ^k^*.

#### 3.1.3 Asymptotic size of the universal set: binary alphabet

By the symmetry of the definition of the mapping ber of *k*-mers mapping function and of the wedges, the same number of *k*-mers map to each wedge, therefore 
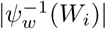
 is independent of *i*, and because the wedges are disjoint, 
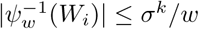
. All the hyperplanes are contained in *W̅*_0_, and therefore 
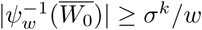
. Hence,

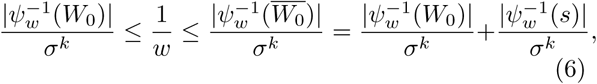

and it suffices to show that the proportion of *k*-mers mapping into *s*, i.e., 
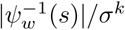
, is asymptotically 0.

Let’s first assume that the alphabet is binary, *σ* = 2.

##### Proposition 3.

The slabs are asymptotically negligible:

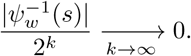

###### Proof.

The number of ways to sum up *n* binary numbers to a value *v* is 
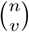
. Hence, the number of *k*-mers mapping to 
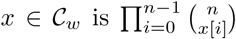
. The point at the center of the cube 
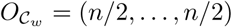
, which is also in all the slabs, has the largest number of *k*-mers mapping to it: 
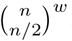
. The volume of a slab, which is bounded by an hyperplane and of thickness *σ* = 2, is *O*(*n^w−1^*).

The images of the mapping *ψ_w_* are the points with integer coordinates, and these points are evenly distributed through out *_w_* 
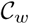
. Hence, the number of points in a slab is proportional to its volume. We can therefore get an over-estimation of the number of *k*-mers mapping into the slabs by estimating the number of *k*-mers mapping to any point in the slab asHence, the proportion of *k*-mers mapping in the slab as the 
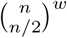
 (the maximum possible), and then multiply by the volume. Unfortunately, the approximation of the proportion of *k*-mers in the *w* − 1 slabs of 
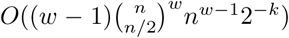
 is too crude and does not converge to 0 when *k* (hence *n*) goes to infinity.

Instead, we split the slabs into 2 parts: (I) the points within a small hypercube 
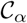
 centered at 
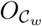
 and of side length 2(*n*/2)*^α^*; (II) the points outside of that hypercube.As we shall see, choosing 1/2 < *α* < *w*/2(*w* − 1) gives the desired convergence. We use 
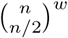
 as an upper-bound on the number of *k*-mers mapping to the points in 
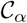
. For the points mapping outside of 
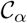
, i.e. in the complement 
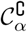
, at least one coordinate is at distance ≥ (*n*/2)*^α^* from *n*/2. The maximum number of *k*-mers mapping to a point *x* of 
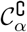
 is when all the coordinates are equal to *n*/2, except for one equal to *n* − (*n*/2)*^α^* or *n* + (*n*/2)*^α^*. Because of the symmetry of the binomial coefficient around *n*/2, 
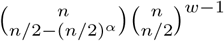
 is an upper-bound on the number of *k*-mers mapping to a point outside of 
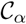
.

In the following *O* expressions, *w* and *σ* = 2 are constants and will be included in the constant in the *O*, while *k* and *n* = *k*/*w* go to infinity.

For (I), from the Stirling approximation 
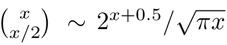
, and the volume of the intersection of 
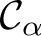
 with the slabs of *O*(*w*(*n*/2)^*α*(*w*−1)^), we have:

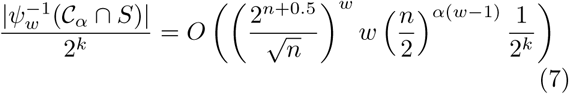

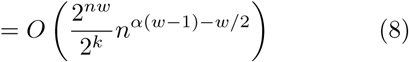

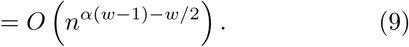

In equation 8, recall that *nw* = *k*. Therefore, 
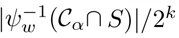
 converges to 0 if the power of *n* in equation 9 is negative, that is if 
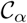
 is not too large and *α* < *w*/2(*w* − 1).

For (II), also from Stirling approximation, we have the equivalence

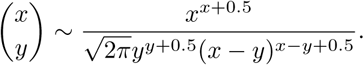

Let *n′* = *n*/2, then *x* = *n* and *y* = *n′* − *n′*^*α*^. Then, the denominator of the binomial coefficient is (except for 
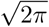
 factor):

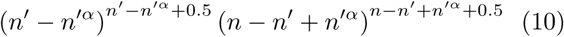

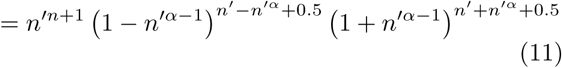

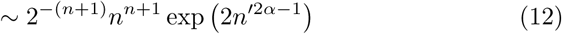

Hence, the proportion of k-mers mapping in the slab outside of 
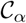
 is:

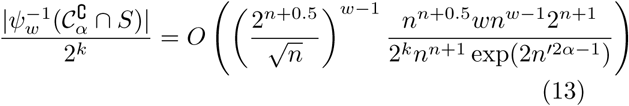

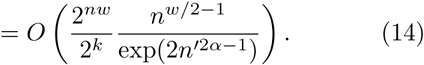

In equation 14, both *n* = *k*/*w* and *n′* = *n*/2 grow linearly with *k*. Therefore, the proportion 
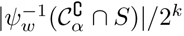
 converges to 0 when the power of *n′* in the exponential is positive, that is when 
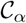
 is large enough and *α* > 1/2.

The following proposition is then a direct consequence of Proposition 3 and Lemma 1

##### Proposition 4.

On the binary alphabet, for any fixed *w*, there exists of sequence of universal sets *U_k_* asymptotically of optimal size, and a sequence of minimizers schemes *f_k_* asymptotically of optimal density.

#### 3.1.4 Asymptotic size of the universal set: even alphabet

We now extend the previous result to even alphabets. Let’s assume that the alphabet Σ is even: |Σ| = *σ* = 2*σ′*. We construct a graph homomorphism (Lichiardopol, 2006; Lempel, 1970) from the de Bruijn graph *D_k,σ_* of order *k* on the alphabet of size *σ* onto the de Bruijn graph *D_k,2_* of order *k* on the binary alphabet. Consider the function *g*(*x*) = ⌊*x*/*σ′*⌋. In other words, *g* maps the first half of the alphabet Σ to 0 and the second half to 1. Then, consider *ϕ*: *D_k,σ_* → *D_k,2_* which applies *g* to each base of a *k*-mer by *σ′*. That is, *y* = *ϕ*(*x*) implies that *y*[*i*] = *g*(*x*[*i*]),*i* ∈ [0,*k* − 1].

It is simple to check that *ϕ* is an onto function and a graph homomorphism: if (*m*, *m′*) is an edge of *D_k,σ_*, then so is (*ϕ*(*m*),*ϕ*(*m′*)) in *D_k,2_*. Inductively, if *m_1_*,…, *m_n_* is a path of *D_k,σ_*, then *ϕ*(*m_1_*),…, *ϕ*(*m_n_*) is a path of *D_k,2_*. Therefore, if *U_w,2_* is a universal set in*D_k,2_* that intersects every path of *w* vertices, then the set *U_w,σ_* = *ϕ*^−1^(*U_w,2_*) is also a universal set of *D_k,σ_*. Moreover, the same number of letters of Σ map to 0 and to 1: |*g^−1^*(0)| = |*g^−1^*(1)| = *σ′*. Then the number of *k*-mers of *D_k,σ_* that map to a *k*-mer *m* of *D_k,2_* is

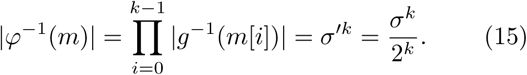

We can now prove a generalization of Proposition 4.

##### Theorem 1.

On an even alphabet, for any fixed *w*, there exists a sequence of universal sets *U_k_* asymptotically of optimal size, and a sequence of minimizers schemes *f_k_* asymptotically of optimal density.

###### Proof.

Let *U_w,2_*(*k*) be the sequence of universal sets of *D_k,2_* constructed in the proof of Proposition 4. Then *U_w,σ_* (*k*) = *ϕ^−1^*(*U_w,2_*(*k*)) is a sequence of universal sets of *D_k,σ_* where |*U_w,σ_*(*k*)| = |*U_w,2_*(*k*)| *σ^k^*/2*^k^*. Therefore, by Proposition 3:

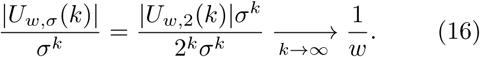

Lemma 1 proves the second part of the statement.

### 3.2 Minimizers asymptotic behavior in *w*

We now consider the converse problem where the length of the *k*-mer is fixed and *w* grows to infinity.

#### Proposition 5.

For any minimizers scheme *f* and any fixed *k*, the density function *w* → *d_f,k,w_* is nonincreasing.

##### Proof.

Let *S_w+k+1_* be any de Bruijn sequence of order *w* + *k* + 1. Let *p_w_* = *S_f,k,w_* (*S_w+k+1_*) and *p_w+1_* = *S_f,k,w+1_*(*S_w+k+1_*) be the set of the positions of the minimizers in *S_w+1+k_* when computing the minimizers for *w* and *w* + 1 respectively. Because the order of the de Bruijn sequence is large enough, *d_f,k,w_* = |*p_w_*|/*σ^w+1+k^* and *d_f,k,w+1_* = |*p_w+1_*|/*σ^w+1+k^*. We now show that *p_w+1_* is a subset of *p_w_*.

Let *ℓ* = *w* +*k* −1. Consider the windows *S_w+k+1_*[*i*, *ℓ*] and *S_w+k+1_*[*i*, *ℓ* + 1], containing *w* and *w* + 1 *k*-mers respectively, both starting at base *i* in *S_w+1+k_*. *S_w+k+1_*[*i*, *ℓ* + 1] contains one extra *k*-mer compared to *S_w+k+1_*[*i*, *ℓ*], the right-most *k*-mer starting at position *i* + *w*. Hence, if a different minimizer is selected in these two windows when computing minimizers for *w* and *w* + 1 respectively, then the *k*-mer at position *i*+*w* must compare less than any *k*-mer in *S_w+k+1_*[*i*, *ℓ*]. Then the *k*-mer at position *i* + *w* also compares less than any *k*-mer in *S_w+k+1_*[*i* + 1, *ℓ*], and the position *i* + *w* is also in *p_w_*.

This previous proposition and the previously known lower-bound on the density, such as 2/(*w* + 1), might suggests that the density of a minimizers scheme goes to 0 asymptotically in *w*. That is not the case however, as shown in this next proposition.

#### Proposition 6.

For any minimizers scheme *f, d_f,k,w_* ≥ *σ*^−*k*^.

##### Proof.

Let *μ* be the *k*-mer that is the lowest for the ordering *f*. In the minimizers scheme, every instance of *μ* in the sequence is the left-most smallest *k*-mer for some window. Hence the algorithm selects as minimizers every instance of *μ*. In a de Bruijn sequence of order *w* + *k*, every *k*-mer occurs the same number of times, *σ^w^* times. Hence the density is ≥ *σ^w^*/*σ^w+k^* = *σ^−k^*.

As the consequence, the expected density factor is not a constant but rather grows at least linearly ≥ (*w* + 1)/*σ^k^*. Moreover this lower-bound is tight.

#### Theorem 2.

For any minimizers scheme *f*, the density *d_f,k,w_* converges asymptotically in *w* to *σ*^−*k*^:

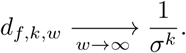

##### Proof.

Let *S_k+w_* be a de Bruijn sequence of order *k* + *w* and *μ* the smallest *k*-mer for the ordering *f*. In every window that contains *μ*, it is the selected minimizer. Let *A_w_* be the set of all the windows of *S_k+w_* that do not contain *μ*. As a worst case scenario, assume that in every window of *A_w_*, a different minimizer is selected. In that worst case, the set of selected positions contains all the *σ^w^* instances of *μ* in *S_k+w_*, and one position in each of the windows of *A_w_*. This gives us an upperbound on the density:

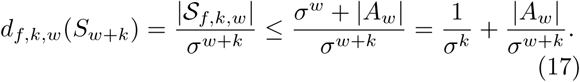

We will show that as *w* increases, the proportion |*A_w_*|/*σ^w+k^* goes to zero, in other words most windows contain *μ* and the only *k*-mer that matters with respect to the density is *μ*.

Because *S_w+k_* is a de Bruijn sequence, every possible window of *w* consecutive *k*-mers occurs exactly *σ* times in *S_w+k_*. Hence, the proportion of interest |*A_w_*|/*σ^w+k^* is exactly the probability of the event that *μ* is not in a window, and we can use a probabilistic argument. The event that a window *ω* does not contain the *k*-mer *μ* is a subset of the event that the non-overlapping *k*-mers starting at positions *i* • *k*, *i* ∈ [0: ⌊*w*/*k*⌋] are not equal to *μ*. Hence,

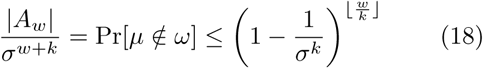

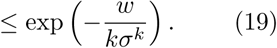

For a fixed *k*, as *w* goes to infinity, this proportion goes to 0. Equation 17 combined with proposition 6 gives the desired limit for *d_f,k,w_*, for all minimizers scheme *f*.

Theorem 2 shows that for very large *w*, all the minimizers schemes are equivalent. Intuitively, when *w* is very large, say *w* » *σ^k^*, every window of *w* consecutive *k*-mers is expected to contain every possible *k*-mer. Hence almost every window contains *μ*, the absolute lowest *k*-mer, and almost no other *k*-mer but *μ* is selected as a minimizer. This happens for any order on the *k*-mers.

For minimizers schemes, the density factor is *θ*(*w*), that is it grows linearly. This does not apply for local and forward schemes as the next proposition exhibits a forward scheme whose density factor is *O*(
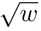
), and therefore a density whose limit is 0 at infinity. The insight for that proof is that a minimizers scheme for parameters *k′*, *w′*, different from *k*, *w*, is a valid local scheme for parameters *k*, *w*, so long as *w′* ≤ *w* and *k′* + *w′* ≤ *w* + *k*. This holds because any function taking a string of length at most *w* + *k* − 1 and returns a value within [0, *w* − 1] is a valid local scheme.

#### Proposition 7.

There exists a forward scheme whose density factor is *O*(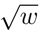).

##### Proof.

Let’s set 
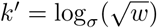
 and *w′* = *w*+*k* − *k′* and consider a minimizers scheme *f′* for parameters *k′*, *w′*. For large values of *w*, *w′* < *w*, and *f′* returns an offset in [0, *w′* − 1] ⊂ [0, *w* − 1], hence *f′* is a valid forward scheme for parameters *k*, *w*. In a de Bruijn sequence of order *w* + *k*, the minimum *k*-mer *μ′* of the ordering *f′* occurs *σ^w+k−k^* times, hence, following the same proof as Theorem 2, the density of *f′* on *S^w+k^* satisfies

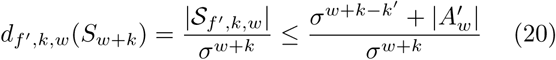

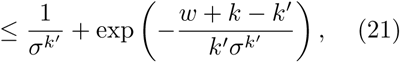

where 
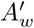
 is the set of windows not containing *μ′*. Consequently, the density factor satisfies

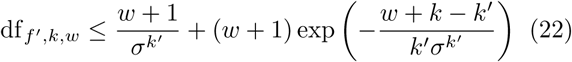

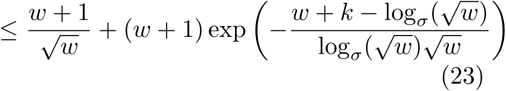

Asymptotically, expression 23 is *O*(
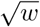
).

Forward schemes do not necessarily have a linear growth for the density factor. The lowest asymptotic density factor achievable by local schemes and forward schemes is still an open question.

### 3.3 Lower bound on forward schemes

When introducing the winnowing scheme, Schleimer *et al*. (2003) provided a lower-bound on the density factor for local schemes of 1.5+1/(2*w*). Unfortunately, their definition of a local scheme differs slightly from ours as they assume that the input *k*-mers are first hashed into fingerprints and that those fingerprints can be assumed independent and uniformly distributed. This is a weaker setting than what is considered here.

Moreover, their theorem does not apply to all local schemes, but only to forward schemes. For a forward schemes, instead of counting the number of selected *k*-mers, we use instead *charged windows*. A window is charged if it is the window with the smallest starting position where a given *k*-mer is selected. More precisely, for a forward scheme *f*, a window is charged if its selected *k*-mer is different than in the preceding window, i.e the window *ω′* starting at position *i* is charged when *i*+*f* (*ω′*) > *i* 1+*f* (*ω_i−1_*). For a forward scheme, the number of charged window is equal to the number of selected *k*-mers and the density is equivalently computed from the number of charged windows. This property, which is not satisfied by general local schemes, is explicitly used in the proof of Schleimer *et al*. (2003) and in the proof of the following theorem.

The theorem below is a refinement of Theorem 1 of Schleimer *et al*. (2003) and it uses the same proof technique. The idea is to look at two windows that are disjoint, and therefore the choice of a selected *k*-mer in each window is independent from the choice in the other window, and to estimate the number of *k*-mers that must be selected between the two windows. We first have a Lemma.

#### Lemma 2.

Let *X* and *Y* be two discrete random variables with values in {0,…, *w* − 1} which are independent and have the same distribution. Then, Pr[*X* ≥ *Y*] ≥ 1/2 + 1/(2*w*).

##### Proof.

Because *X* and *Y* are IID, Pr[*X* > *Y*] = Pr[*Y* > *X*], hence

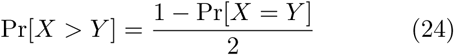

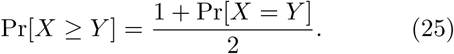

We now get a lower bound on Pr[*X* = *Y*]. Let 
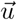
 be a vector whose coordinates are Pr[*X* = *i*] for *i* ∈ {0,…, *w* − 1}, and let 
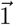
 be the vector of all 1s. Because the variables are IID, 
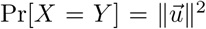
, and by the Cauchy-Schwartz inequality

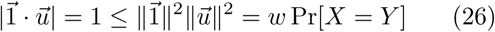

#### Theorem 3.

The density of any forward scheme satisfies

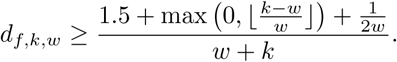

##### Proof.

Let *f*: Σ*^w+k−1^*[0, *w* − 1] be a forward scheme, and consider two windows of *w* consecutive *k*-mers, *ω_1_* = *S*[*i*, *w* + *k* − 1] and *ω_2_* = *S*[*i* + *w* + *k* + 1, *w* + *k* − 1], starting respectively at base *i* and *i* + *w* + *k* + 1 (see Figure 2). The last base of *ω_1_* is at index *i* + *w* + *k* − 2, hence there is no shared sequence between the two windows, and a gap of 2 bases between the two windows. Without loss of generality, we can assume that the input sequence is a de Bruijn sequence of large enough order so that every possible pair of windows *ω_1_* and *ω_2_* is encountered, the same number of times. We can therefore use a probabilistic argument: the starting position *i* is chosen at random, consequently the strings *ω_1_* and *ω_2_* are IID.

We are counting the number of charged windows starting at positions in the interval [*i*+1, *i*+*w*+*k*], that is all the windows after *ω_1_* and before *ω_2_*. Consider the random variables of the offset of the selected *k*-mer in each window, *X_1_* = *f* (*ω_1_*) and *X_2_* = *f* (*ω_2_*). Because the windows *ω_1_* and *ω_2_* do not share any sequence, *X_1_* and *X_2_* are independent and have the same distribution. The respective positions of the selected *k*-mer in *ω_1_* and *ω_2_* is *s_1_* = *i* + *X_1_* and *s_2_* = *i* + *w* + *k* + 1 + *X_2_*.

At least one window with starting index in [*i*+1, *s_1_* + 1] must be charged, as the *k*-mer selected in window *ω_1_* is not in the window starting at position *s_1_* + 1. Moreover, additional windows must be charged if the number of bases between *s_1_* and *s_2_* (excluding bases at *s_1_* and *s_2_*) is larger or equal to 2*w*. If Δ is the distance between *s_1_* and *s_2_*, the number of extra *k*-mers to select is ⌊(Δ − *w*)/*w*⌋ if Δ ≥ *w*, 0 otherwise. The number of bases between these two selected *k*-mers is

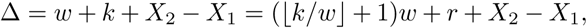

where *k* = ⌊*k*/*w*⌋*w* + *r*. Given that both *X_1_* and *X_2_* are in [0, *w* − 1] and *r* ≥ 0, *X_2_* − *X_1_* + *r* ≥ −*w* + 1. Therefore Δ ⌊*k*/*w*⌋*w* and at least an extra ⌊(*k* − *w*)/*w*⌋ windows must be charged (provided that *k* ≥ *w*).

When Δ ≥ (⌊*k*/*w*⌋ + 1)*w*, or equivalently when *X_2_* ≥ *X_1_* − *r*, at least one extra window is charged. By Lemma 2, this event occurs with probability

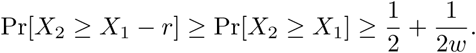

Hence, in any interval [*i* + 1, *i* + *w* + *k*] of length *w* + *k*, the expected number of charged windows is at least 1.5 + max(0, ⌊(*k* − *w*)/*w*⌋) + 1/(2*w*), and the density of charged windows is as stated.

**Figure 2:**
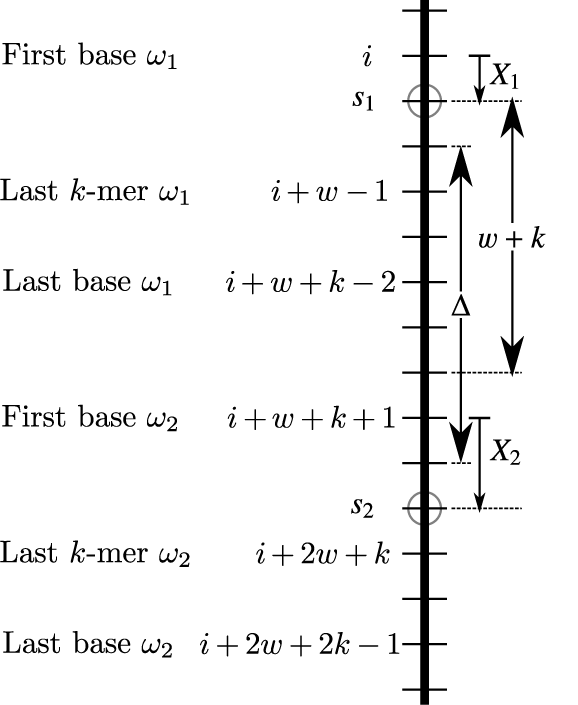
The sequence is by a thick vertical, and each base is a tick mark. In this example, *k* = 3, *w* = 4, *X_1_* = *f* (*ω_1_*) = 1 and *X_2_* = *f* (*ω_2_*) = 2. The first base, last *k*-mer start and last base of *ω_1_* and *ω_2_* are *i*, *i* + *w* − 1, *i* + *w* + *k* − 2 and *i* + *w* + *k* + 1, *i* + 2*w* + *k*, *i* + 2*w* + 2*k* − 1 respectively. The selected *k*-mers are marked with a circle at positions *s_1_* and *s_2_*. ∆ is the number of bases between, and excluding, the selected *k*-mers. There are *w* + *k* windows that can be charged after *ω_1_* and before *ω_2_*, between bases *i*+1 and *i*+*w*+*k* included.

This lower-bound is tight for some extremal cases. For *w* = 1, the lower-bound on the density is 1, which is the value of the density factor for any scheme. When *k* → ∞, the lower-bound on the density goes to 1/*w*, which is achieved asymptotically by the orderings constructed in Section 3.1.

On the other hand, when *w* → ∞, the lower-bound on density factor goes to 1.5. It is still an open question whether this bound can be reached asymptotically by a forward scheme.

## 4 Discussion

### 4.1 Asymptotic behavior in *w*

Figure 3 summarizes the known upper and lowerbounds for local, minimizers and forward schemes, for a fix parameter *k* and varying *w*.

**Figure 3:**
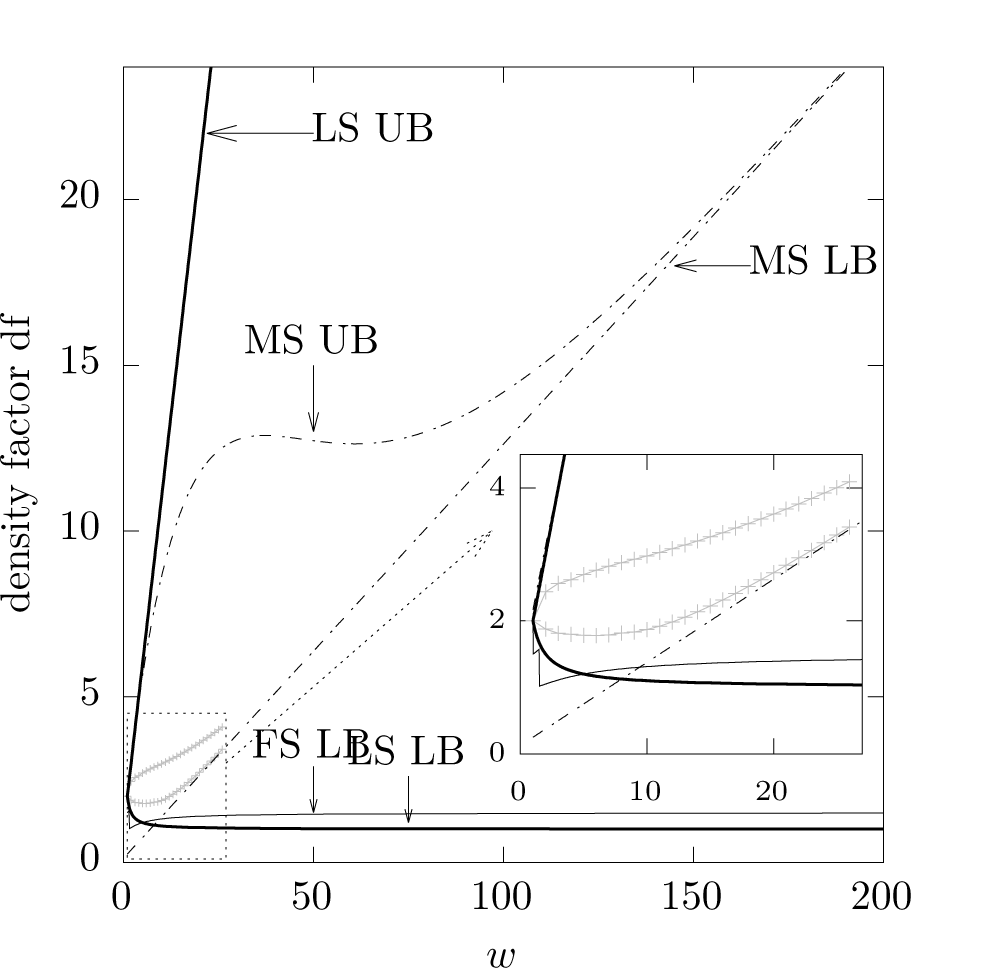
For a binary alphabet *σ* = 2 and a fix value of *k* = 3, this plots show the upper-bound and lowerbound for the local-schemes (thick lines “LS UB” and “LS LB”) and for the minimizer-schemes (dashed lines “MS UB” and “MS LB”). The thin line “FS LB” is the lower-bound for forward scheme. In addition, in the inset, the gray lines show the actual best and worse density factor achievable by a minimizer schemes for *k* = 3 and for *w* ∈ [1, 27].

The dashed lines show the upper and lower-bound for minimizers schemes. In addition, we computed through exhaustive search (there are “only” 2^3^! = 40 320 different orderings) the actual lowest and highest density factor achievable for *k* = 3 and *w* ∈ [1, 27]. These values are shown in gray, and the inset zooms in on that region of the graph. The minimizers scheme is the most understood scheme, and the known bounds are tight asymptotically in *w*. Theorem 2 shows that for very large *w*, all the minimizers schemes are equivalent, as the number of selected positions is dominated by the occurrences of the lowest *k*-mer *μ*, and this number is the same for any ordering. This is responsible for the linear lower-bound on the density factors. Local and forward schemes do not have that inherent limitation (see Proposition 7).

The thicker lines show the upper and lower-bounds for a local schemes. The upper-bound is tight, as the constant function *f* (*ω*) = 0, that always picks the first *k*-mer in any window, selects every *k*-mer in the sequence and therefore has a density factor of *w* + 1. On the other hand, it is not known if the trivial lowerbound for local schemes (i.e. one *k*-mer per window hence a density factor of 1 + 1/*w*) is tight or not, even for asymptotic *w*. Theorem 3 shows that this trivial lower-bound cannot be achieved by a forward scheme (the thin line).

#### Local schemes are more powerful

It would be conceivable that for any set of parameters *k*, *w*, there is always a forward scheme that is among the local schemes with the lowest density. Then, the lowerbound of Theorem 3 would also apply to local schemes. However, this is not the case. By formulating the problem of finding a local scheme with lowest density as an Integer Linear Program (ILP), we found set of parameters (e.g. *k* = 2 and *w* = 4) where none of the lowest density solutions are forward schemes. Therefore, the lower-bound on forward schemes may not apply to local scheme in general.

The local schemes is the largest class of schemes and the least understood. In fact, most of what is known about local schemes is derived from our knowledge of the minimizers schemes and forward schemes. The previous remark shows that the local schemes are strictly more powerful than the other type of schemes, and that the lower-bounds on the density that were previously thought to constrain local schemes (say *d* ≥ 1.5/(*w* + 1)) may not apply. New insights and a deeper understanding of local schemes are necessary to design local schemes with even lower densities. For example, to achieve a low density, a local scheme must not be a forward scheme, that is it must sometimes do backward jumps (i.e. pick *k*-mers in a non-increasing manner). Why such backward jumps are beneficial to reduce the number of selected *k*-mers is not yet understood.

### 4.2 Asymptotic behavior in *k*

Figure 4 shows, for varying values of *w*, the density obtained by orderings compatible with the universal sets constructed in section 3.1. The orderings are constructed such that *k*-mers mapping into the wedge *W_0_* compare less to the *k*-mers mapping into the slabs, who also compare less than the *k*-mers mapping outside of *W_0_*. Ties are resolved using lexicographic order.

**Figure 4:**
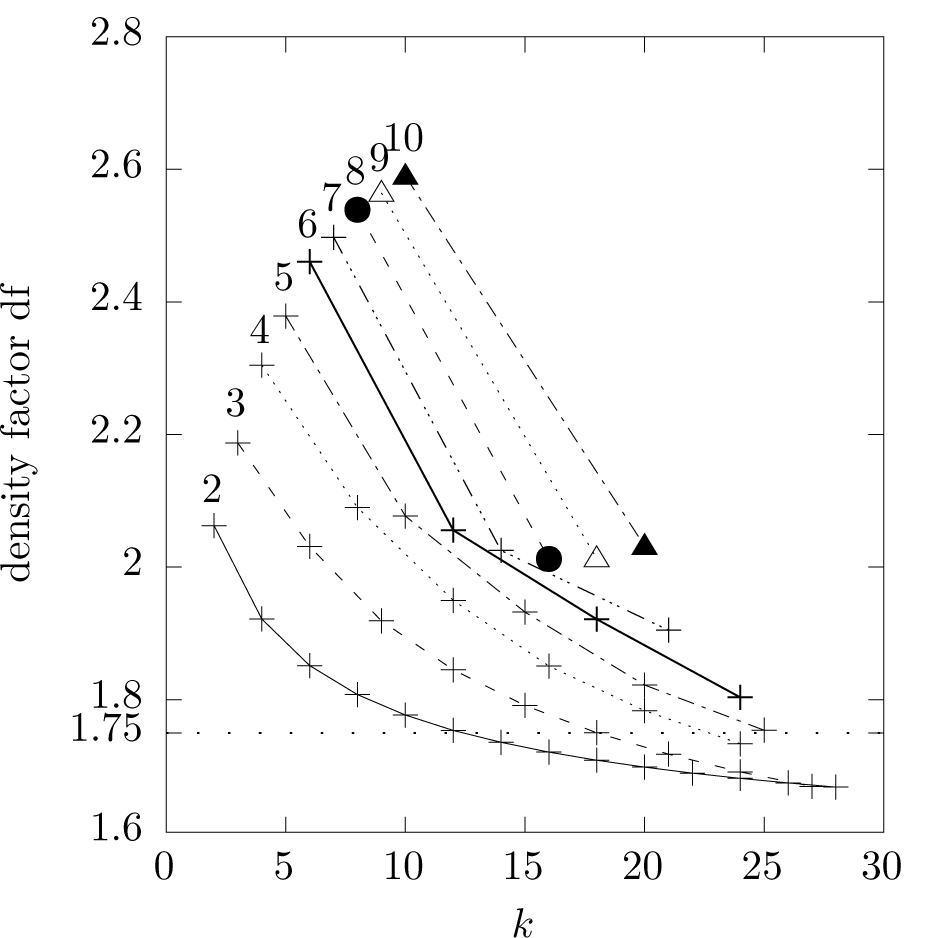
Each curve represents, for a fixed value of *w*, the density factor as a function of *k* obtained by minimizers schemes compatible with the universal sets constructed in section 3.1. The value of *w* is written above the top left most point of each curve. The horizontal dotted line shows the previously believed lower bound for *w* = 2.

For *w* = 2, and *k* ≥ 14, the density factor obtained is below 1.5 + 1/(2*w*) = 1.75, that is below what was previously thought possible. Moreover, such a range of values for the parameters *k* and *w* is potentially useful for some applications. According to Theorem 1, all of the curves on Figure 4 have an asymptote of 1 + 1/*w*, although such low density factors may be reached for relatively large values of *k*, larger than useful in practice. Better orderings compatible with the universal sets than the simple one used here might exist to achieve lower density for smaller values of *k*. Also, even though local schemes are not asymptotically in *k* more powerful than minimizers schemes, local schemes might achieve lower densities for smaller values of *k* and therefore be more practical than the order used here.

One benefit of the construction of the universal sets in section 3.1 is that it can be implemented as a test. That is, there is an indicator function using *O*(*k*) memory and *O*(*k*) time to check if a *k*-mer is in the universal set: for a given *k*-mer *m*, computing the vector *ψ_w_* (*m*) requires *k* additions. There is no need to pre-compute the universal set or hold it in memory. Similarly, the comparison between two *k*-mers takes *O*(*k*) time.

#### Minimum size universal sets

In Orenstein *et al.* (2017), we proposed the problem of finding universal sets of minimum size in the de Bruijn graph, and gave a heuristic algorithm to find small, although not necessarily minimum, universal sets. In particular, the use of a heuristic was justified by the difficulty of finding vertex cover for path of length *ℓ* in a Directed Acyclic Graph (DAG) (Paindavoine and Vialla, 2015). Although this problem is NP-hard for general DAGs, the construction given in section 3.1 provides a solution that is asymptotically optimal in the particular case of de Bruijn graphs.

#### Cycle structure of the de Bruijn graph

In the following we propose a high level and intuitive description of why the construction given in section 3.1 works asymptotically, when *k* is much larger than *w*. Consider a cycle *C_=_* of length *ℓ* = *w* in *D_k_*. This cycle could have one node in each of the wedges *W_i_*. On the other hand, a smaller cycle *C_<_* of length *ℓ* < *w* cannot have a node in each of the *w* wedges. Therefore, it must have some nodes, if not all, inside of the slabs, close to the diagonal of the hypercube 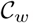. For a larger cycle *C_>_* of length *ℓ* > *w*, if *ℓ* is a multiple of *w*, the cycle could rotate around the wedges *W′*, with no nodes falling in the slabs. If *ℓ* is not a multiple of *w*, some nodes of *C_>_*, but not all, must fall in the slabs as well.

Let 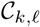 be the number of cycles of length *ℓ* in *D_k_*. For a fixed *ℓ*, the function 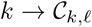 is increasing until *k* = *ℓ* − 1, then it is constant (Maurer, 1992). In other words, when *k* becomes large, the number of cycles of length ≤ *w* remains constant, while the number of larger cycles grows, and it grows very quickly. Asymptotically, we can “ignore” the fine grain cycle structure of the de Bruijn graph as the behavior is dominated by long cycles, which tend to be more regular with respect to our embedding.

Conversely, for values of *k* in the same order of magnitude as *w*, the precise cycle structure of the de Bruijn graph matters, and designing small universal sets or low density schemes requires taking this cycle structure into account.

## 5 Conclusion

In this study, through the asymptotic analysis of minimizers, forward and local schemes, we deepened the theoretical understanding of these techniques, and thereby showed that greater improvements than previously thought are possible. In particular, we completely characterized the behavior asymptotically in *k* and gave an efficient algorithm to create the first known optimal minimizers schemes. Because the minimizers schemes are the weakest type of schemes, this shows that all schemes are optimal asymptotically in *k*. For forward schemes, we gave a refined lower-bound that applies to all parameters *k* and *w*.

The asymptotic behavior in *w* is markedly different than the asymptotic behavior in *k*. For large *w*, the lower-bound for the minimizers schemes is higher than for the forward scheme, which is higher than for local schemes.

The local schemes are not well understood at all. Although we do have some examples of optimal local schemes found through ILP or brute force search, there is currently no algorithm to generate local schemes with low density. Every algorithm proposed so far is a forward scheme. A greater understanding of local schemes holds, at least for large values of *w*, the greatest promise to design even better schemes.

